# Geometric control of frequency modulation of cAMP oscillations due to Ca^2+^-bursts in dendritic spines

**DOI:** 10.1101/520643

**Authors:** D. Ohadi, P. Rangamani

## Abstract

The spatiotemporal regulation of cAMP and its dynamic interactions with other second messengers such as calcium are critical features of signaling specificity required for neuronal development and connectivity. cAMP is known to contribute to long-term potentiation and memory formation by controlling the formation and regulation of dendritic spines. Despite the recent advances in biosensing techniques for monitoring spatiotemporal cAMP dynamics, the underlying molecular mechanisms that attribute to the subcellular modulation of cAMP remain unknown. In the present work, we model the spatio-temporal dynamics of calcium-induced cAMP signaling pathway in dendritic spines. Using a 3D reaction-diffusion model, we investigate the effect of different spatial characteristics of cAMP dynamics that may be responsible for subcellular regulation of cAMP concentrations. Our model predicts that the volume-to-surface ratio of the spine, regulated through the spine head size, spine neck size, and the presence of physical barriers (spine apparatus) is an important regulator of cAMP dynamics. Furthermore, localization of the enzymes responsible for the synthesis and degradation of cAMP in different compartments also modulates the oscillatory patterns of cAMP through exponential relationships. Our findings shed light on the significance of complex geometric and localization relationships for cAMP dynamics in dendritic spines.

## Introduction

Dendritic spines, small bulbous protrusions from the dendrites of neurons are the main excitatory synaptic sites that compartmentalize postsynaptic responses. Spine dynamics are intimately associated with longterm potentiation (LTP), long-term depression (LTD), and synaptic plasticity [1,2]. The influx of calcium due to neurotransmitter-release and the associated gating of ion channels is universally accepted as the first step towards these processes [3,4]. However, spines are more than hotbeds of electrical activity; recent studies have shown that dendritic spines are subcompartments of signaling and biochemical activity downstream of calcium influx [5,6] and there is a tight coupling between electrical and chemical activity in spines [7]. In particular, the connection between calcium dynamics and cAMP/PKA activation is one of the key elements for connecting the short-time scale events associated with calcium influx to the longer time scale of structural plasticity [8–10].

In response to calcium influx, cAMP transients have been reported in neurons [11] and cAMP/PKA dynamics are tightly coupled to that of calcium [12–14]. In a companion study, we developed a computational model for calcium-induced cAMP/PKA activity in neurons [15] and predicted that the cAMP/PKA pathway acts as a leaky integrator of calcium signals. We also experimentally showed that calcium spontaneously oscillates in dendritic spines of hippocampal subregional volumes, Cornu Ammonis 1 (CA1) neurons [15]. Another key factor that regulates LTP is the geometry of dendritic spines [16–18]. Spine morphology can affect synaptic potential integration in dendrites [19] and their variety of shapes and sizes provide high functional diversity [20]. The head shape, neck length, and neck diameter in spines can change during the synaptic plasticity [21]; changes in spine morphology and spine density are associated with learning and memory [21]. In other cell types, it is well-known that the spatial localization of cAMP regulating molecules and spatial aspects of signaling can govern its dynamics [22,23]. Two key types of enzymes that produce and degrade cAMP are believed to be responsible for cAMP subcellular compartmentalization. Phosphodiesterases are suggested to act as a sink and a diffusion barrier and form cAMP microdomains [24–26]. Additionally, adenylyl cyclases can be colocalized with other components of the pathway and generate cAMP locally [12,27]. However, little is known about the molecular mechanisms of such compartmentalization.

These observations led us to the following questions: can geometric and spatial features of spines regulate the periodically forced cAMP/PKA oscillations due to oscillating calcium dynamics? If so, how? To answer these questions, we developed a three-dimensional reaction-diffusion model of calcium-induced cAMP/PKA dynamics in dendritic spines and focused on how two critical spatial aspects – spine geometry and enzyme localization – change cAMP/PKA dynamics. We studied the effect of features that regulate volume-to-surface ratio; spine head size and neck size along with the presence or absence of the spine apparatus as a physical barrier for molecular diffusion. Because we know that many of these enzymes are not uniformly distributed, we also investigated the role of localization of AC1 and PDE4. Our results show that spatial localization of enzymes on the spine membrane and in the cytoplasm can alter the temporal response of cAMP to induced calcium transients. Thus, in addition to kinetic properties as shown previously [15], we predict that the spatial organization of molecules in the dendritic spine can affect how cAMP/PKA dynamics respond to calcium input.

## Model Development

### Model assumptions

In order to simulate the spatiotemporal dynamics of cAMP dendritic spines, we developed a reaction-diffusion model that accounts for the different biochemical species (see Table S1 and Table S2 of Supplemental Information for the list of reactions and the list of parameters) and their localization on the plasma membrane or in the cytosol and boundary fluxes (Figure 1). We briefly discuss the assumptions made in developing this model and outline the key equations below.

**Figure 1:**
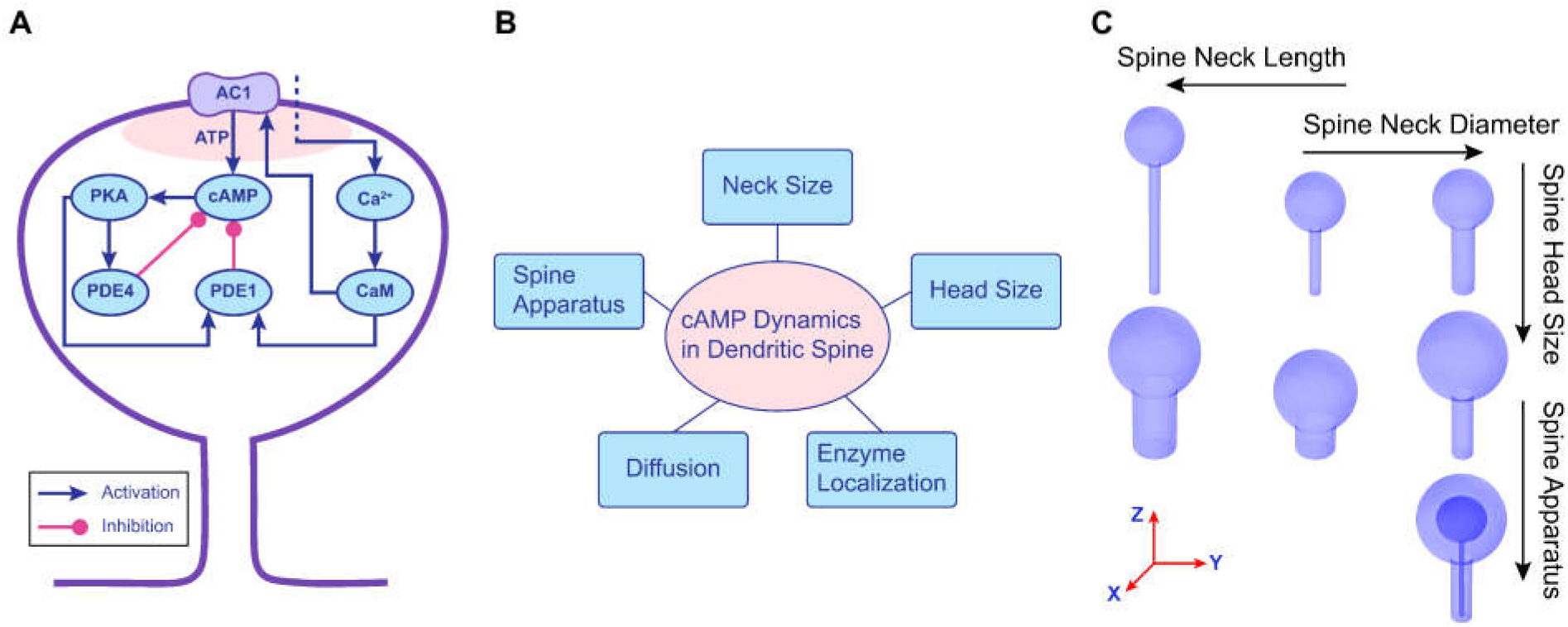
cAMP pathway in dendritic spines. (A) A schematic of the spatially organized reaction network for the cAMP-PKA pathway in dendritic spines. (B) In this study, we consider the different geometric and spatial aspects that affect cAMP dynamics in dendritic spines. (C) 3D geometries of the spines used in this model to study cAMP dynamics; these geometries are designed to study the role of different spine head size, neck sizes and the presence and absence of the spine apparatus.

- **Time scales:** We focus on cAMP/PKA dynamics at time scales of tens of minutes. Calcium dynamics are modeled as sinusoidal oscillations with exponential decays based on the experimental observations on the second and minute timescales. The second-scale oscillations (0.5 Hz) were set up based on experimental observations in our companion study [15], and minute-scale oscillations (0.003 Hz) were inspired by Gorbunova and Spitzer’s observations [11].
- **Spine head:** Spine volumes range from 0.003 to 0.55 *μm*^3^, spine neck diameters are within a range of 0.04 to 0.5 *μm*, and the total length of spines are between 0.2 and 2 *μm* [28] in the hippocampal CA1 region. Based on the head and neck shape, spines are classified as stubby, thin (<0.6 *μm* in diameter) and mushroom spines (>0.6 *μm* in diameter) [28]. Here, we study two spherical heads with two different spine head volumes: 0.065 *μm*^3^ and 0.268 *μm*^3^, which are within the range of experimentally measured spine head volumes (Figure 1C).
- **Spine neck:** The spine neck is modeled as a cylinder with a diameter of 0.1, 0.2, and 0.4 *μm* and length of 1.32, 0.66, and 0.33 *μm* representing a thin, an average, and a thick neck, respectively. Thin spines are known as small head spines with a thin and long neck. Mushroom spines are spines with a large mushroom shaped head with a thick and short neck. The geometric specifications of the studied spine geometries are shown in Table 1 and in Figure 1C.
- **Postsynaptic density:** Postsynaptic density area in hippocampal CA1 region ranges from 0.008 to 0.54 *μm*^2^ [28]. We chose the size of the postsynaptic density (PSD) based on the correlation between the head volume and PSD area reported by Allerano et al. [29] to localize cAMP synthesis on the spine head surface. Calcium enters the spine through NMDA glutamate receptors influx. Postsynaptic calcium is released by presynaptic glutamate binding to NMDA-type glutamate receptor and removal of Mg^2+^ block as a result of postsynaptic depolarization [30]. The input calcium function (see details in section S4 of Supplemental Information) enters the spine head from PSD area (Figure 1A). The resting cytosolic calcium is 0.1 *μm* and it can rise up to 1 *μm* [31]. Calcium oscillations for the model have been designed based on the concentration range and timescale of calcium oscillations in neurons [11,15].
- **Spine apparatus:** In order to investigate the effect of physical barriers such as spine apparatus, we modeled large spines with spine apparatus. The size of the spine apparatus is a spheroid head with a=0.225, b=0.225, c=0.200 *μm* and a cylindrical neck with D=0.05 *μm* and L=0.823 *μm* (*V_SA_* =0.044 *μm*^3^) (Table 1d). The size of the spine apparatus is taken from image constructions of the spines by Wu et al. [32] (Table 1d). In this model, the membrane of the spine apparatus acts as a reflective barrier for cAMP/PKA without any sink or source terms.
- **Plasma membrane fluxes:** In our model, AC1, AC1· Ca_2_· CaM, and AC1·Ca_4_·CaM are localized on the plasma membrane while all other species are in the spine volume. We do not explicitly include the various calcium channels and pumps on the PSD but rather prescribe the calcium profile in the spine.
- **Deterministic approach:** We assume that all molecules in the model have large molecular concentration and model the reaction-diffusion equations (RDEs) using deterministic approaches.

**Table 1:** Geometric specifications of the different spines considered in this study

| Spine Name | Table 1a: Spines with different head sizes |  |  |  |  |  |
| --- | --- | --- | --- | --- | --- | --- |
|  | Head Diameter | Neck Diameter | Neck Length | Spine Surface | Spine Volume | Volume/Surface |
| Control | $0.5 \mu m$ | $0.2 \mu m$ | $0.66 \mu m$ | $1.174 \mu m^2$ | $0.086 \mu m^3$ | $0.073 \mu m$ |
| L.Avg.Avg* | $0.8 \mu m$ | $0.2 \mu m$ | $0.66 \mu m$ | $2.400 \mu m^2$ | $0.288 \mu m^3$ | $0.120 \mu m$ |
|  | Table 1b: Spines with different neck sizes |  |  |  |  |  |
|  | Head Diameter | Neck Diameter | Neck Length | Spine Surface | Spine Volume | Volume/Surface |
| S.Tn.Ln | $0.5 \mu m$ | $0.1 \mu m$ | $1.32 \mu m$ | $1.194 \mu m^2$ | $0.076 \mu m^3$ | $0.064 \mu m$ |
| S.Tn.Avg | $0.5 \mu m$ | $0.1 \mu m$ | $0.66 \mu m$ | $0.986 \mu m^2$ | $0.070 \mu m^3$ | $0.071 \mu m$ |
| Control | $0.5 \mu m$ | $0.2 \mu m$ | $0.66 \mu m$ | $1.174 \mu m^2$ | $0.086 \mu m^3$ | $0.073 \mu m$ |
| L.Avg.Avg | $0.8 \mu m$ | $0.2 \mu m$ | $0.66 \mu m$ | $2.400 \mu m^2$ | $0.288 \mu m^3$ | $0.120 \mu m$ |
| L.Tc.Avg | $0.8 \mu m$ | $0.4 \mu m$ | $0.66 \mu m$ | $2.732 \mu m^2$ | $0.347 \mu m^3$ | $0.127 \mu m$ |
| L.Tc.Sh | $0.8 \mu m$ | $0.4 \mu m$ | $0.33 \mu m$ | $2.317 \mu m^2$ | $0.305 \mu m^3$ | $0.132 \mu m$ |
|  | Table 1c: Spines with or without spine apparatus |  |  |  |  |  |
|  | Head Diameter | Neck Diameter | Neck Length | Spine Surface | Spine Volume | Volume/Surface |
| L.Avg.Avg | $0.8 \mu m$ | $0.2 \mu m$ | $0.66 \mu m$ | $2.400 \mu m^2$ | $0.288 \mu m^3$ | $0.120 \mu m$ |
| L.Avg.Avg.Sa | $0.8 \mu m$ | $0.2 \mu m$ | $0.66 \mu m$ | $2.400 \mu m^2$ | $0.244 \mu m^3$ | $0.102 \mu m$ |
| | | | | | $V_{SA}=0.044 \mu m^3$ | |
|  | Table 1d: Spine apparatus geometry |  |  |  |  |  |
|  | SA Head Shape | SA Head Size | SA Neck Diameter | SA Neck Length | SA Volume | SA Surface |
| | Spheroid | a=$0.225 \mu m$<br>b=$0.225 \mu m$<br>c=$0.200 \mu m$ | $0.05 \mu m$ | $0.823 \mu m$ | $0.044 \mu m^3$ | $0.719 \mu m^2$ |

\*The first letter in the spine name represents the size of the spine head (S: small; L: large). The second part of the spine name represents its neck diameter (Tn: thin; Avg: average; Tc: thick). The third part of the spine name represents its neck length (Ln: long; Avg: average; Sh: short). The fourth part of spine name represents the spine apparatus (Sa: with spine apparatus). Spine names without the fourth part do not have a spine apparatus.

Based on these assumptions, we constructed a 3-dimensional spatial model of calcium-induced cAMP/PKA pathway in dendritic spines. Our control geometry is a small sized spine with an average neck and a total volume of ~0.086 *μm*^3^ without a spine apparatus. While all the simulations are conducted in 3D (Figure 1C), results from the simulations are shown for a 2D cross-section for simplicity and ease of interpretation.

### Governing equations

The spatiotemporal dynamics of each species, *c*, in the volume is given by a reaction-diffusion equation as

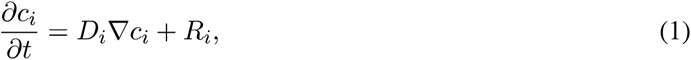

where *c_i_*, *i* ∈ {1, 2,…, 21}, represents the concentration of the *i^th^* species as a function of time and space, *D_i_* is the diffusion coefficient, ∇ represents the Laplacian operator in 3D, and *R_i_* is the net reaction flux for the *i^th^* species. The diffusion coefficients of different species and the initial concentration of the different species are shown in Table S3. For membrane-bound species, the volume concentrations are converted to surface concentrations by multiplying them by the volume-to-surface ratio.

### Boundary condition at the PM

Flux boundary conditions that balance diffusive flux with reaction rate are used to represent reactions that take place at the plasma membrane between molecules on the membrane and in the volume. There are four species for which these boundary conditions apply. Specifically, in this case, Ca^2+^ and Ca_2_·CaM binding with AC1 and the enzymatic conversion of ATP to cAMP due to the action of PM-bound AC1 becomes a time-dependent flux boundary condition for both cAMP and ATP, which can be written as

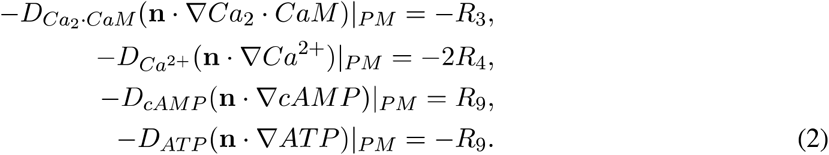

Here, n represents the normal to the surface. These time-dependent fluxes in the boundary conditions closely couple the temporal responses encoded in the reaction terms and the curvature response encoded in the normal vector in the diffusive flux term [33].

### Boundary condition at the spine apparatus membrane

We assume that the spine apparatus is purely a diffusive barrier and does not play an active role in modulating any of the biochemical dynamics. Therefore, for all the species, the boundary condition at the spine apparatus is a Neumann boundary condition, given as

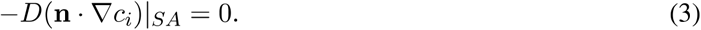

### Geometries used in the model

We modeled the dendritic spines using simplified geometries of spheres. Dendritic spines consist of a spine head attached to a neck, with a similarly structured spine apparatus within the spine (Figure 1C). The different geometries used in the model are shown in Table 1.

### Numerical Methods

Simulations were conducted using the commercially available finite-element software COMSOL Multiphysics 5.3 [34]. In order to solve our system of partial differential equations, we used time-dependent general partial differential equations and general boundary partial differential equations modules [34]. Starting with a coarse and unstructured mesh, we decreased the mesh size until we obtained the same results when using the maximum mesh size. COMSOL was allowed to optimize the element sizes through the “physics-controlled mesh” option. The linear system was solved directly by using the MUMPS solver. Newton’s method (nonlinear method) was used to linearize the system. Time integration was performed using a backward differentiation formula (BDF) with both adaptive order and adaptive step sizes. The mesh statistics are shown in Table 3.

**Table 2:**
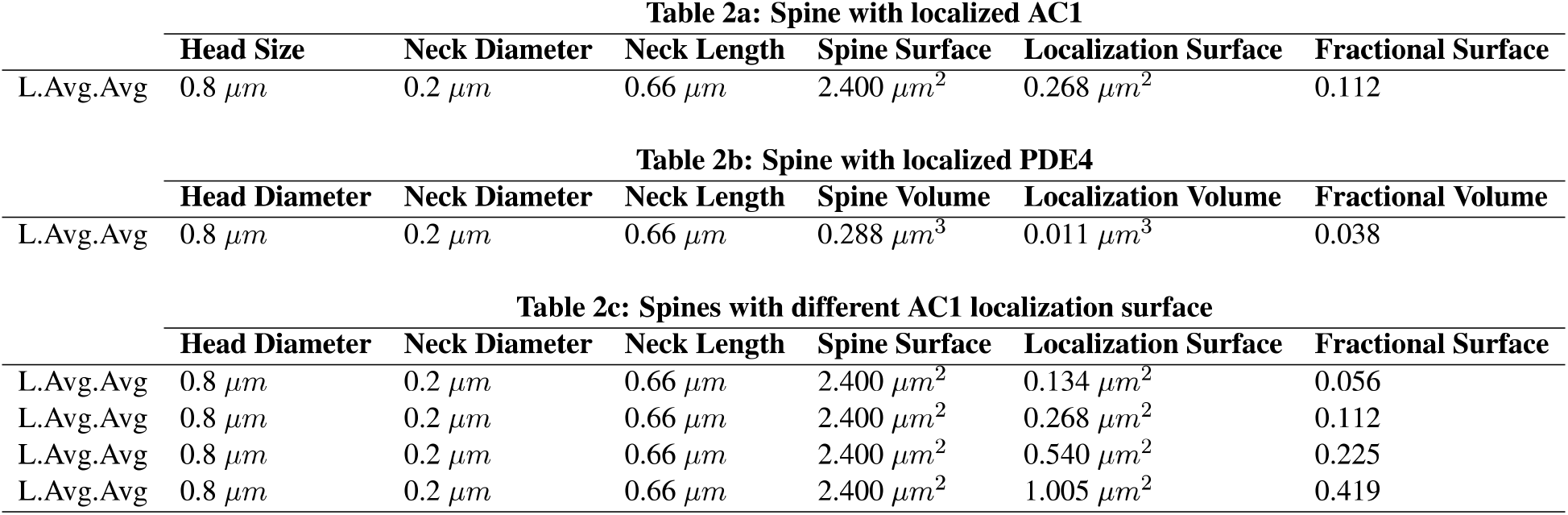
Parameters for fractional area localization of AC1

**Table 3:** Mesh statistics for the control spine (spherical spine with small head and average neck)

| Element type | Number of elements | Min element quality | Avg element quality | Element volume ratio | Mesh volume |
| --- | --- | --- | --- | --- | --- |
| Tetrahedron | 6006 | 0.2229 | 0.6423 | 4.921E-4 | 8.32E-20 $m^3$ |
| Element type | Number of elements | Min element quality | Avg element quality | Element area ratio | Mesh face area |
| Triangle | 1638 | 0.3982 | 0.8263 | 0.004638 | 1.177E-12 $m^2$ |
| Element type | Number of elements | Element length ratio | Mesh edge length |  |  |
| Edge Element | 278 | 0.1238 | 9.944E-6 $m$ | | |
| Element type | Number of elements |  |  |  |  |
| Vertex Element | 31 |  |  |  |  |

### Metrics for cAMP/PKA dynamics

In order to compare cAMP concentration across different spine geometries shown in Table 1, we normalized the cAMP concentrations in each case with respect to the cAMP concentration in the control spine to compare across different geometries. We multiplied cAMP concentration by 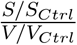, in which, *S_Ctrl_* and *V_Ctrl_* are total surface area and total volume of the control geometry and S and V are the total surface area and the total volume of the geometry under consideration. In addition to studying the spatiotemporal dynamics of cAMP in spines of different geometries, we also compare the peak time for each period, the peak concentration, and the Area Under the Curve (AUC) for each period. These metrics allow us to compare the oscillatory behaviors of cAMP in dendritic spines [35,36].

## Results

### Spatiotemporal dynamics of cAMP/PKA in dendritic spines

We studied the spatiotemporal dynamics of cAMP and PKA in response to calcium influx in a dendritic spine. The calcium input and the resulting cAMP dynamics in the control spine are shown in Figure 2. The calcium stimulation patterns are based on a sinusoidal function with 0.5 Hz pulses and 5-minute separation between bursts and an exponential decay along each burst [15]. In-phase oscillations of calcium and cAMP have been reported in neurons [11] and other cell types [37] and the oscillation timescales are based on the suggested oscillation period for cAMP and calcium in the literature [11,38]. We observe that while there is no discernible gradient of calcium or cAMP in these spines, the temporal pattern of oscillations demonstrates the shift in peak time through the progression of the pathway. For instance, while calcium concentration peaks at 601 seconds (10 min), activated AC1 peaks at 615 seconds and activated PDE1 peaks at 607 seconds (Figure 2B). As a result, cAMP concentration oscillates with the larger timescale as shown previously and peak concentration is at 656 seconds (11 min) (Figure 2B).

**Figure 2:**
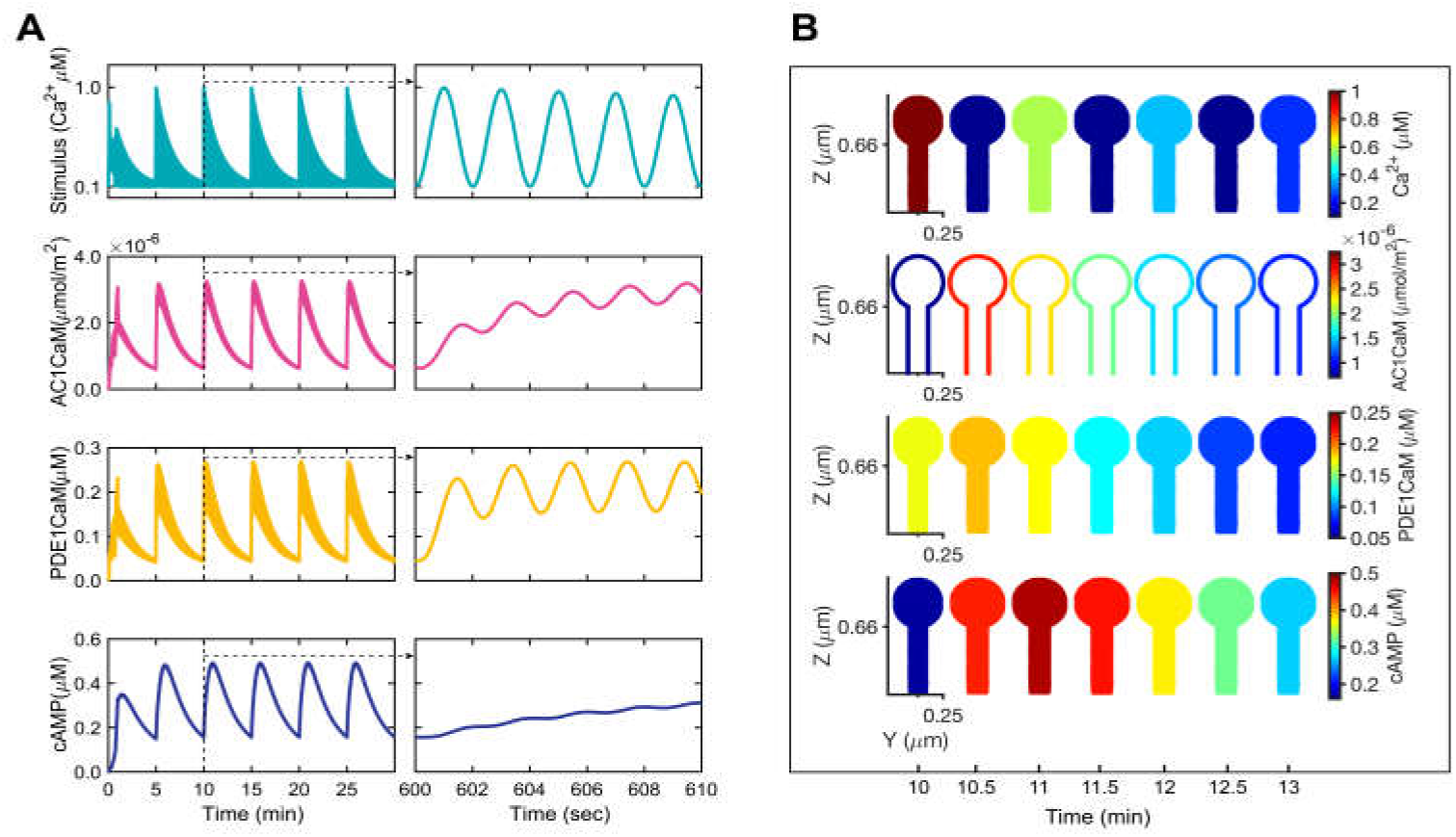
Oscillatory dynamics of Ca^2+^, AC1·CaM, PDE1·CaM, and cAMP in the spatial model. (A) Ca^2+^, AC1·CaM, PDE1·CaM, and cAMP dynamics in a 3D control spine. The stimulus is a calcium input with a frequency of 0.5 Hz and five-minute bursts, oscillating between 0.1 *μm* (calcium at rest) and 1 *μm* (max calcium concentration). The figure insets show that the concentration profiles on the second-scale (in the range of 600 to 610 seconds). (B) Ca^2+^, AC1·CaM, PDE1·CaM, and cAMP spatial maps during one burst of calcium (one oscillation) show a variation in concentration corresponding to the amplitude in (A) but no spatial gradients. While AC1·CaM is a membrane-bound species on the surface, all other species shown here are in the volume.

### cAMP dynamics are affected by modulating spine volume-to-surface ratio

The volume-to-surface area ratio is an important characteristic of spine geometry and organization because it accounts for the effect of both volume and membrane reaction fluxes. Volume-to-surface ratio can be modified in multiple ways by changing the spine head size, spine neck size, and by the presence or absence of a spine apparatus. We investigated the effect of volume-to-surface ratio on cAMP dynamics in our model by systematically varying these geometric features (Table 1).

#### Effect of spine head size

In order to study the effect of the spine head size, we considered two spherical heads with different sizes (D_1_ =0.5 *μm* and D_2_ =0.8 *μm*), while maintaining the same neck diameter and neck length (Table 1a) resulting in different volume-to-surface ratios. We observed that for the same calcium input, the head size plays a significant role in defining the oscillatory pattern of cAMP dynamics (Figure 3A). These temporal patterns map to the spatial patterns shown in Figure 3B. As the volume-to-surface area ratio of the spine decreases, cAMP concentration increases, as expected. A spine with a smaller head, which has lower volume-to-surface ratio shows not just a higher, but also a broader range of concentrations from the peak to the base (from t=11 to 15 min). The cAMP concentration in all different cases in this study is normalized with respect to the control spine. No discernible difference was observed between cAMP concentrations at different points inside a given geometry. Furthermore, we noticed that the change in spine head size not just changed the concentration of cAMP, but also altered the oscillatory dynamics as characterized by the peak time, AUC, and the peak amplitude (Figure 3C). Increasing the volume-to-surface ratio of the spine by modulating the spine head size results in lower peak amplitude of cAMP concentration, lower area under the curve, and a delay in the peak time. This is because the change in the volume-to-surface area ratio affects the time-dependent flux boundary conditions shown in Equation (2).

**Figure 3:**
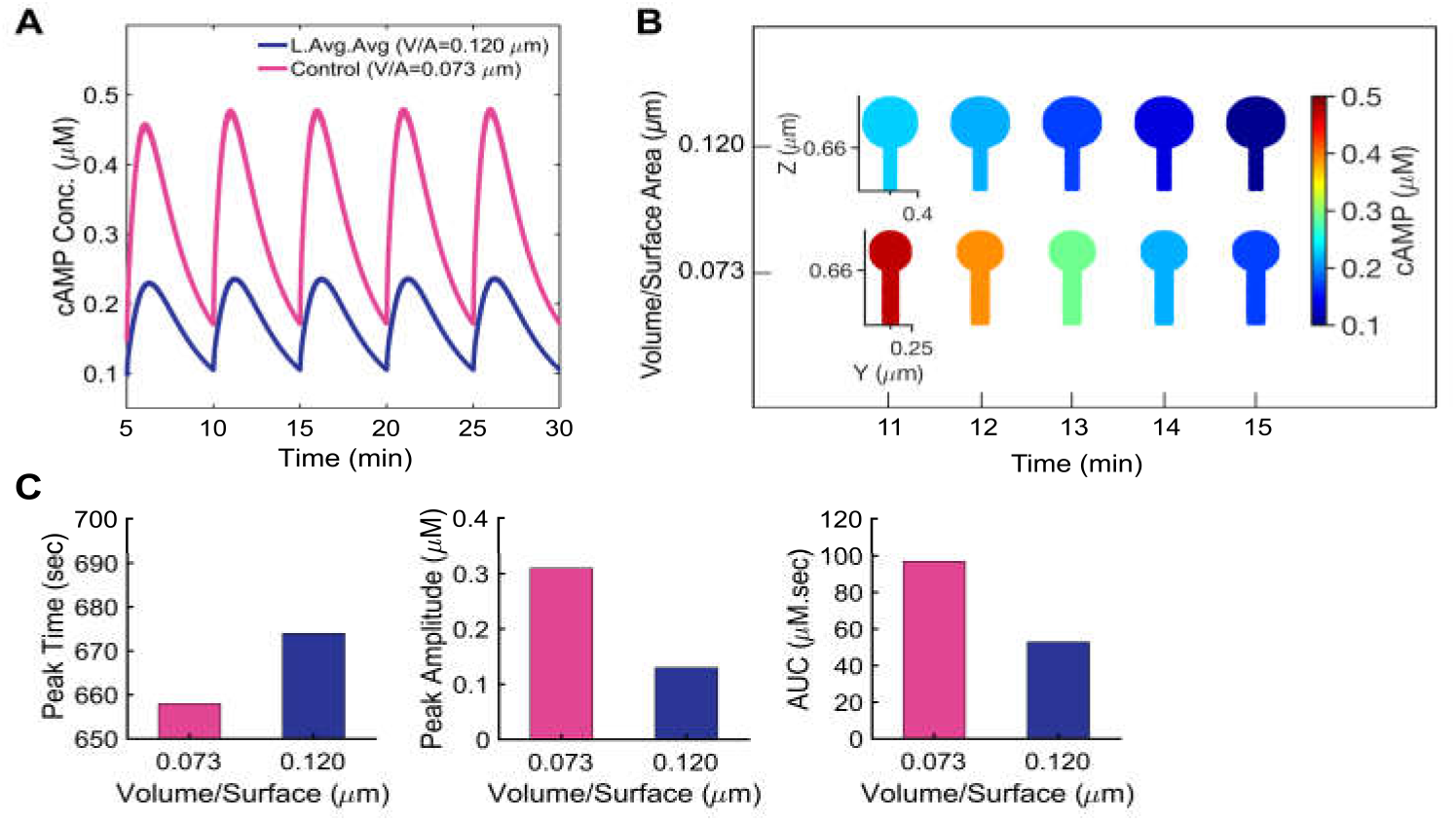
Effect of the spine head size on the cAMP dynamics. (A) Effect of spine head size on the cAMP concentration profile. (B) Spatial maps of cAMP concentration profile in spines with different head sizes shown in (A) during one oscillation of cAMP show that decreasing the volume-to-surface area can increase the cAMP concentration substantially. (C) Effect of spine head size on the peak time (second peak shown in A), peak amplitude, and area under the curve (during one oscillation period). The spine with larger head shows a lower peak amplitude, a lower area under the curve, and a (16-second) delay in the peak time compared to that of the smaller spine, reflecting the coupling between boundary conditions due to membrane-bound species.

#### Effect of spine neck size

Small, thin spines (D<0.6 *μm*) are usually associated with a thin and long neck and mushroom spines (D>0.6 *μm*) usually have a thicker and shorter neck in comparison to thin spines [39]. We investigated how changes to spine neck size, which affect the volume-to-surface ratio of spines affect cAMP dynamics (Figure 4). We constructed geometries that reflect different combinations of spine head and neck sizes (Table 1b). These neck sizes are in the range of experimentally measured spines for each of these two thin and mushroom categories [29,39]. Decreasing the neck diameter from 0.2 to 0.1 *μm* and increasing the neck length from 0.66 to 1.32 *μm* increases the cAMP concentration (Figure 4A). In other words, by decreasing the volume-to-surface area ratio of the spines, the cAMP concentration increases (Figure 4B) and the spine with small neck diameter and long neck length shows the highest peak amplitude, the highest area under the curve, and earliest peak time (Figure 4E). For mushroom-like spines, increasing the neck length from 0.33 to 0.66 *μm* and decreasing the neck diameter from 0.4 to 0.2 *μm*, the cAMP concentration in spines with large heads increases (Figure 4C). Similar to small head spines, in spines with large heads by decreasing the volume-to-surface ratio, cAMP concentration increases (Figure 4D) and the thinnest and the longest neck shows the highest peak amplitude and area under the curve with earliest peak time (Figure 4E).

**Figure 4:**
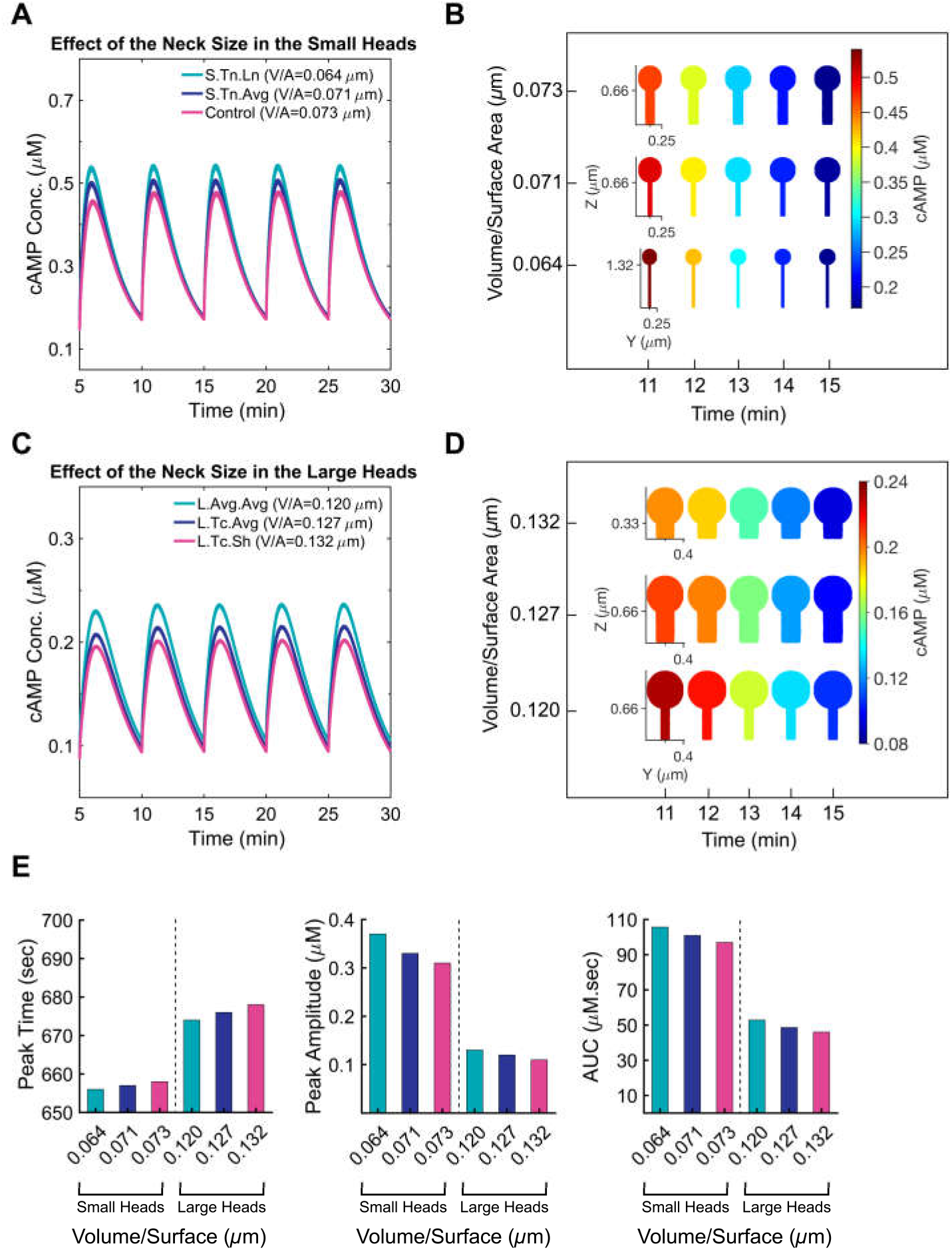
Effect of neck size on cAMP dynamics in spines with different spherical head sizes. (A) Comparison of cAMP concentration in three spines with the same head size (D=0.5 *μm*) but different neck diameters (0.1 and 0.2 *μm*) and different neck lengths (0.66 and 1.32 *μm*). (B) Spatial maps of cAMP concentration in spines with a small head (D=0.5 *μm*) and different neck sizes during a five-minute oscillation period. (C) cAMP concentration in three spines with larger heads in comparison to those shown in (A) (D=0.8 *μm*) with different neck lengths (0.66 and 0.33 *μm*), and different neck diameters (0.2 and 0.4 *μm*). (D) Spatial maps of cAMP in spines with large heads (D=0.8 *μm*) during one oscillation period. (E) The spine with the highest volume-to-surface area ratio has the lowest peak amplitude, the lowest area under the curve, and the highest delay in the peak time.

#### Effect of spine apparatus

An additional geometric feature of dendritic spines is their internal organization; roughly more than 80% of the large mushroom spines in hippocampal CA1 dendrites of adult rats have a specialized endoplasmic reticulum called the spine apparatus [40]. To understand the role of internal organelles such as spine apparatus that can act as a physical barrier, especially in spines with larger heads (D>0.6 *μm*), we modeled the cAMP pathway in a spine with a large head (D=0.8 *μm*) and a spine apparatus. The spine apparatus was modeled as a spheroid with a=0.225, b=0.225, c=0.200 *μm* and a cylindrical neck with D=0.05 *μm* and L=0.823 *μm* (Table 1d). Presence of the spine apparatus decreases the volume-to-surface ratio and thereby increases cAMP concentrations (Figure 5A, B). As a result, we predict that the presence of the spine apparatus as a physical barrier to diffusion decreases the volume-to-surface area and increases the peak amplitude and the area under the curve and expedites the peak time (Figure 5C).

**Figure 5:**
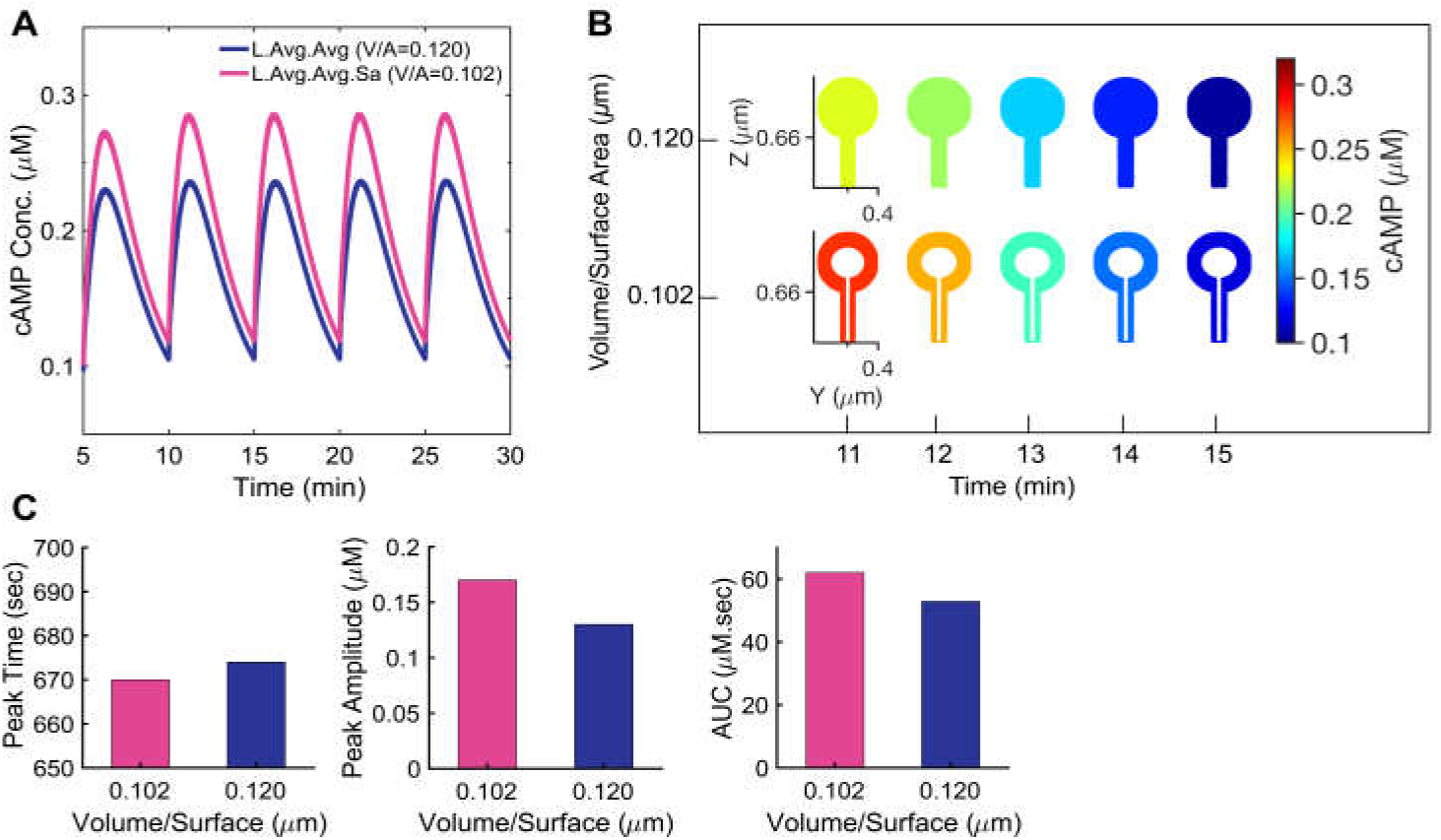
Effect of spine apparatus (SA) as a diffusion physical barrier on cAMP concentration in spines with large heads. (A) Effect of SA on cAMP concentration in large heads (D=0.8 *μm*). (B) cAMP concentration maps of the spine with a large head with and without SA during one oscillation period. (C) Effect of SA on peak time, peak amplitude, and area under the curve. The spine with spine apparatus shows a higher peak amplitude, a higher area under the curve, and its peak time precedes the spine without the SA.

#### cAMP dynamics are modulated by localized synthesis and degradation through enzyme localization

One of the key features of cAMP dynamics is the localization of the cyclase and PDEs [23,25,41]. The coupling between enzyme localization and cAMP microdomains has been hinted at in the literature but has not been explicitly considered in our model so far. To investigate how localization of these molecules affects cAMP dynamics, we considered the following scenarios: localization of membrane-bound molecules (AC1· Ca_2_ · CaM and AC1· CaM) to the head surface; localization of PDE4 in the spine head; and localization of both AC1 and PDE4 (Table 2a, b and Figure 6A). We set the size of the AC1 localization area to be 0.268 *μm*^2^ based on the head volume/PSD area correlation reported by [29]. Interestingly, we observed that by localizing AC1 on the head surface in a large head, the oscillation amplitude of cAMP increases by almost 10-fold (Figure 6B). Localization of PDE inside of the head volume, on the other hand, only has a negligible effect on cAMP concentration. However, localization of both AC1 and PDE further increases the cAMP concentration relative to the non-localized case (Figure 6B). Figure 6C shows the cAMP concentration profile characteristics for non-localized, localized AC1, localized PDE4, and localized AC1 and PDE4. AC1 localization seems to increase the peak amplitude and the area under the curve substantially and cause a significant delay relative to the non-localized case (almost 40 seconds). The spatial maps of AC1 localization show a delay in the peak time occurrence (Figure 6D). However, PDE4 localization with much lower cAMP concentration does not show this delay (Figure 6E).

**Figure 6:**
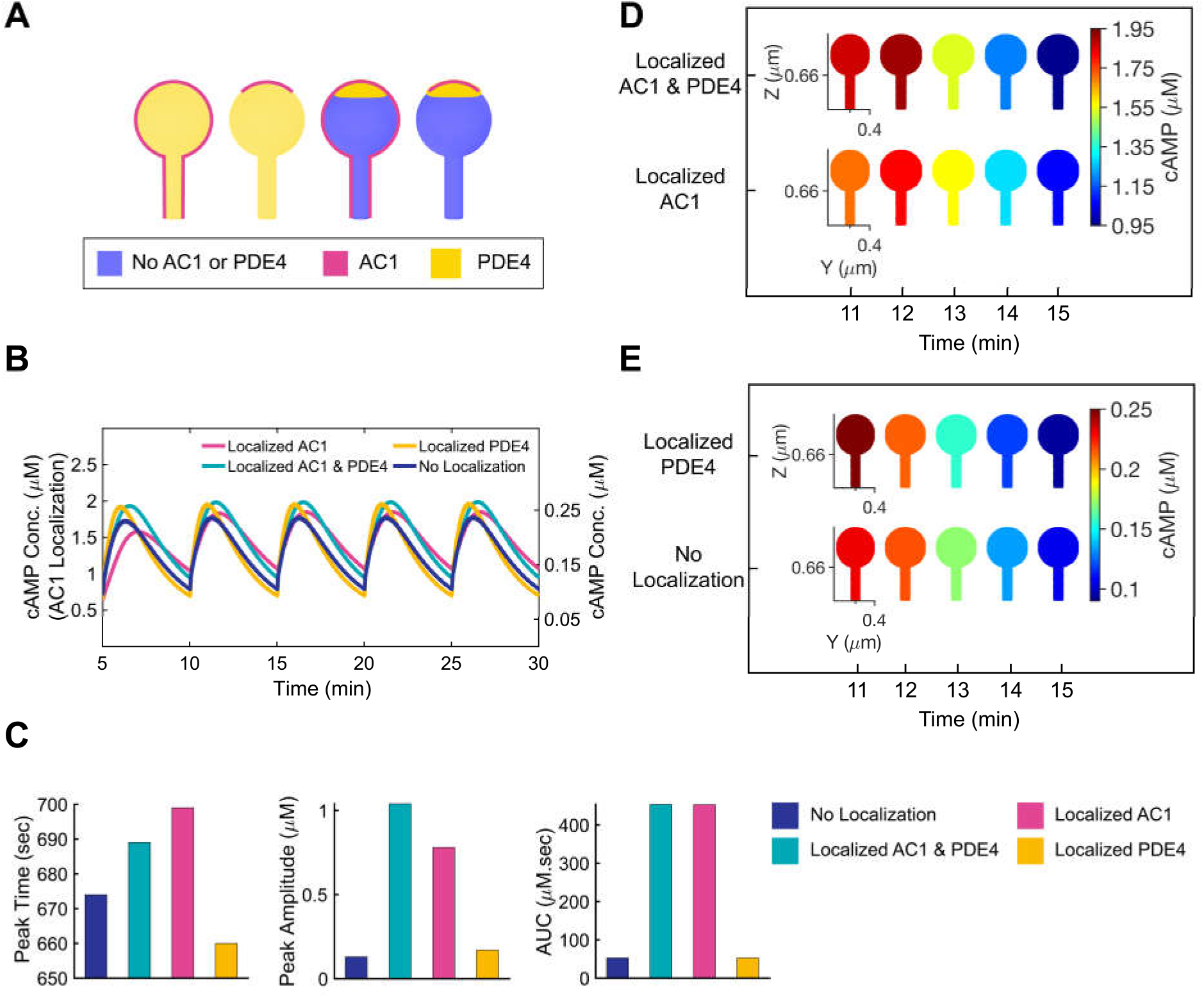
Effect of enzyme localization on cAMP dynamics in dendritic spines with a large head (D=0.8 *μm*) and average neck size (D=0.2 *μm*, L=0.66 *μm*). (A) Schematic of AC1 localization on the spine head surface and PDE4 localization in the spine head volume in the large head. (B) Effect of AC1 and PDE4 localization in the large head on cAMP dynamics. (C) Effect of enzyme localization on the peak time, peak amplitude, and area under the curve for one oscillation period. AC1 localization increases the peak amplitude and area under the curve substantially and causes a delay in the peak time, while PDE4 localization shifts the peak time backward and expedites the peak time. (D) Comparison of spatial maps of AC1 localization and both AC1 and PDE4 localization in the large head. (E) Effect of PDE4 localization on spatial maps of cAMP in the large head.

Since localization of AC1 seems to affect the temporal response of cAMP through membrane fluxes (boundary conditions), we next asked if the fractional area of localization could tune the temporal dynamics in a deterministic matter. In order to investigate how the area of AC1 localization on the spine head can shift the cAMP concentration peak time, peak amplitude, and area under the curve, we studied the AC1 localization effect in four different localization surface areas (Table 2c). We found that decreasing the localization area size shifts the peak time further forward relative to the non-localized case and increases the peak amplitude and the area under the curve (Figure 7A). We observed that the shift in peak time, peak amplitude increase, and area under the curve increase showed an exponential relationship with the fractional area of localization (Figure 7B). These exponential relationships hint at well-defined size-shape relationships between spine size and function.

**Figure 7:**
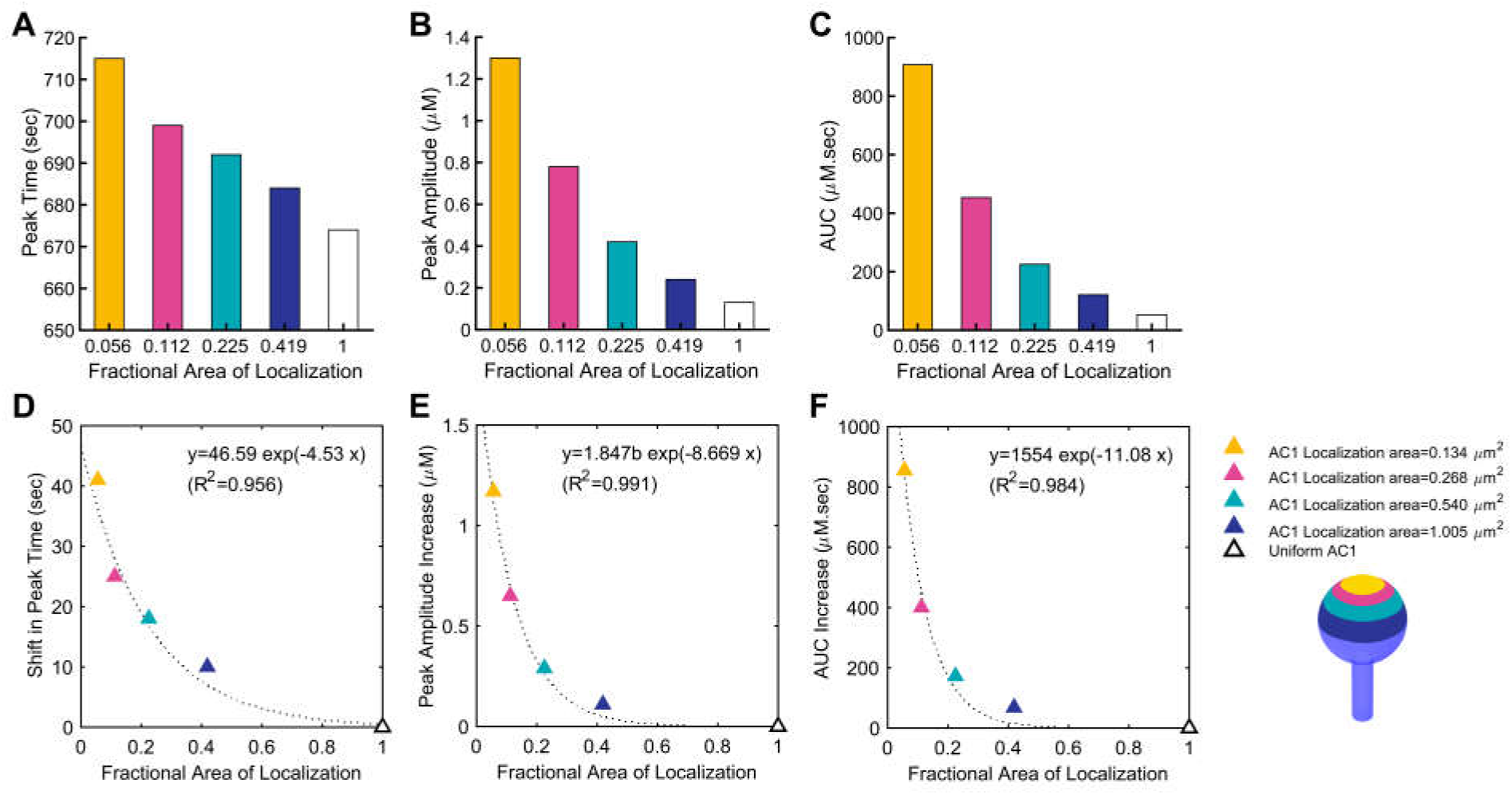
Effect of AC1 localization area on cAMP oscillation pattern: (A) Effect of AC1 localization area on the cAMP peak time for four different AC1 localization areas: 0.134, 0.268, 0.540, and 1.005 *μm*^2^. The fractional area of localization is calculated by dividing the localization area by the total surface area (2.4 *μm*^2^) of a spine with a large head (D=0.8 *μm*) and an average neck (D=0.2 *μm*, L=0.66 *μm*). Decreasing the localization area to 0.134 *μm*^2^ (fractional area=0.056) causes a delay of almost 40 seconds in the peak time. (B) Effect of AC1 localization area on the peak amplitude. Decreasing the localization area to 0.134 *μm*^2^ (fractional area=0.056) shows an almost 10-fold increase in the peak amplitude. (C) Effect of AC1 localization area on the area under the curve (AUC) during one oscillation period. The area under the curve increases substantially (almost 17-fold) by decreasing the AC1 localization fraction to 0.056. (D) Decreasing the AC1 localization area with respect to the total spine surface area shifts the cAMP oscillation peak time forward and in comparison to the uniform AC1 case, the shift in peak time increases exponentially. (E) The cAMP oscillation peak amplitude changes exponentially by changing the AC1 localization area. (C) The area under the curve is another cAMP oscillation property that changes exponentially by changing the AC1 localization area.

## Discussion

In neurons, cAMP oscillations are thought to have an important role in regulating the pulsatile release of hormones such as gonadotropin-releasing hormone [42,43] and axon guidance [44,45]. It is also well-known that calcium spikes are necessary for changes in cAMP concentrations, but only certain bursts of calcium spikes increase cAMP levels in neurons [11]. The dynamics of calcium-induced cAMP has been modeled by us and others [15,22,24,46] with a focus on identifying the mechanisms underlying interdependent oscillations. We showed that cAMP is primarily sensitive to the longer timescale effects of calcium rather than the shorter time scales [15]. Here, we investigated how spatial features of dendritic spines such as spine size and ultrastructure and localization of enzymes can impact the dynamics of cAMP. We expected that spatial aspects of cAMP dynamics would simply reflect the temporal behavior of cAMP as observed in a well-mixed model [15]. However, we found that geometric factors can have unexpected effects on the temporal dynamics of cAMP. Our findings and model predictions can be summarized as follows: *first*, spine volume-to-surface ratio, which can be modulated through spine head size, spine neck geometry, and by the presence or absence of the spine apparatus, affects the temporal dynamics of cAMP. Furthermore, we found that increasing volume-to-surface ratio increases the peak time, decreases the peak amplitude, and decreases the AUC of cAMP exponentially (Figures 3 to 5). ***Second***, spatial localization of cAMP producing enzymes (AC1) and cAMP-degrading enzymes (PDE1) also impact the temporal response of cAMP in response to calcium oscillations (Figure 6). The temporal dynamics of cAMP oscillations depend then not only on the calcium influx, but also on the fractional area of localization of these enzymes (Figure 7).

The first prediction is particularly relevant in the context of spine size and shape variation during development and disease [47-49]. In the adult human hippocampus almost 65% of spines are thin spines (small bulbous shaped head with a diameter of smaller than 0.6 *μm*), 25% are mushroom spines (mushroom shaped head with a diameter of larger than 0.6 *μm*), and the rest are stubby, multisynaptic, filopodial, or branched [39,50]. Furthermore, spine geometry is thought to restrict the diffusion of both cytosolic and membrane-bound molecules [51]. In general, a low volume-to-surface ratio in different parts of a cell has been suggested to be responsible for generating cAMP gradients in finer structures [52–55]. We have found that different ways of modulating the volume-to-surface area ratio alter cAMP response predictably and point to the role of both spine membrane surface area and spine volume. These geometric characteristics are important in considering how the complex geometry of a spine can affect the dynamics of these different molecules.

Other important factors responsible for compartmentalization of cAMP can be colocalization of key components of the pathway by scaffold proteins, such as AKAP79 [56,57]. A-kinase anchoring proteins (AKAPs) are known to tether PKAs to specific sites to phosphorylate GluR1 receptors or facilitate AC action specificity [58–61]. We found that localization of AC1 and PDE1 can substantially change the cAMP concentration level, oscillation amplitude, and peak time (Figure 6) and these features depend on the fractional area of localization (Figure 7).

Our model predictions on geometric regulation of cAMP dynamics in dendritic spines have implications for spatial control of information processing in spines and size-function relationships in structural plasticity [62-64]. We predict that frequency control of cAMP dynamics in response to calcium influx occurs not only through kinetic mechanisms [15], but also due to spatial regulation of volume-to-surface area globally and locally. These results also point towards the need to study the role of local volume-to-surface ratios in realistic geometries such as those developed by Wu *et al.* [32]. Despite our model predictions, there are a few limitations of our work that must be acknowledged. Our model assumes a uniform diffusion coefficient of cAMP, which may not be the case in the crowded environment of the spine head [16,65,66] (see section S6, Figure S3 of Supplemental Information for the effect of diffusion coefficient of cAMP and Ca^2+^). We further assume that ATP is available in large quantities and is not rate limiting (see section S5, Figure S2 of Supplemental Information for ATP dynamics). However, we know that mitochondria are positioned at the base of the dendrite and their size can scale with synaptic plasticity [67-69]. Therefore, the role of ATP availability in the spine head and diffusion of ATP through the neck and the crowded head remains to be explored. Furthermore, in this model, we have assumed that the spine apparatus is simply acting as a diffusion barrier for cAMP. However, in longer time scales, both mitochondria and spine apparatus couple cAMP, calcium, and ATP dynamics and these effects will need to be included. These and other effects are the focus of current and future studies in our group.

## Acknowledgments

The authors wish to thank Dr. Danielle L. Schmitt, Miriam Bell, Justin Laughlin, and Allen Leung for their valuable comments. The authors also thank Dr. Thomas M. Bartol for insightful discussions. This work was supported by AFOSR MURI grant number FA9550-18-1-0051 to PR.

## Author Contributions

DO and PR conceived the study, analyzed the data, and wrote the manuscript; DO performed the simulations.

## Abbreviations

CaM: Calmodulin
AC1: Adenylate cyclase 1
ATP: Adenosine triphosphate
cAMP: Cyclic adenosine monophosphate
PDE1: Phosphodiesterase 1
PDE4: Phosphodiesterase 4
PKA: Protein kinase A
SA: Spine apparatus
ER: Endoplasmic reticulum
AKAP: A-kinase-anchoring protein
AUC: Area under the curve
RDE: Reaction-diffusion equation
Cornu Ammonis 1: CA1

