## Supplemental Information for "Geometric control of frequency modulation of cAMP oscillations due to Ca^2+^-bursts in dendritic spines"

D. Ohadi and P. Rangamani*

Department of Mechanical and Aerospace Engineering, University of California, San Diego, La Jolla, CA 92093

*Corresponding Author, Department of Mechanical Aerospace Engineering, SME 344G, Jacobs School of Engineering, University of California, San Diego La Jolla, CA 92093-0411

### S1 Table of Reactions

TableS 1: cAMP-PKA pathway reactions, reaction types, and reaction rates used in the model

| | List of Reactions | Reaction Type | Reaction Rate |
|---|---|---|---|
| Module 1: | $Ca^{2+}$/CaM Complex Formation | | |
| 1 | $2\,Ca^{2+} + CaM \xrightleftharpoons[kb_1]{kf_1} Ca_2{\cdot}CaM$ | mass action | $R_1 = k_{f1}[Ca^{2+}]^2[CaM] - k_{b1}[Ca_2CaM]$ |
| 2 | $2\,Ca^{2+} + Ca_2{\cdot}CaM \xrightleftharpoons[kb_2]{kf_2} Ca_4{\cdot}CaM$ | mass action | $R_2 = k_{f2}[Ca^{2+}]^2[Ca_2CaM] - k_{b2}[Ca_4CaM]$ |
| Module 2: | Enzyme Activations By CaCaM Complex | | |
| 3 | $AC1 + Ca_2{\cdot}CaM \xrightleftharpoons[kb_3]{kf_3} AC1{\cdot}Ca_2{\cdot}CaM$ | mass action | $R_3 = k_{f3}[AC1][Ca_2CaM] - k_{b3}[AC1Ca_2CaM]$ |
| 4 | $2\,Ca^{2+} + AC1{\cdot}Ca_2{\cdot}CaM \xrightleftharpoons[kb_4]{kf_4} AC1{\cdot}Ca_4{\cdot}CaM$ | cooperative binding | $R_4 = \frac{k_{f4}[AC1Ca_2CaM][Ca^{2+}]}{K_{m4}+[Ca^{2+}]} - k_{b4}[AC1Ca_4CaM]$ |
| 5 | $PDE1 + Ca_2{\cdot}CaM \xrightleftharpoons[kb_5]{kf_5} PDE1{\cdot}Ca_2CaM$ | mass action | $R_5 = k_{f5}[PDE1][Ca_2CaM] - k_{b5}[PDE1Ca_2CaM]$ |
| 6 | $2\,Ca^{2+} + PDE1{\cdot}Ca_2{\cdot}CaM \xrightleftharpoons[kb_6]{kf_6} PDE1{\cdot}Ca_4{\cdot}CaM$ | cooperative binding | $R_6 = \frac{k_{f6}[PDE1Ca_2CaM][Ca^{2+}]}{K_{m6}+[Ca^{2+}]} - k_{b6}[PDE1Ca_4CaM]$ |
| Module 3: | cAMP Production | | |
| 7 | $ATP + \mathbf{AC1{\cdot}Ca_4{\cdot}CaM} \xrightarrow{Kcat_7} cAMP$ | Michaelis-Menten | $R_7 = \frac{K_{cat7}[AC1Ca_4M][ATP]}{K_{m7}+[ATP]} - K_{r7}[cAMP]$ |
| Module 4: | PKA Formation | | |
| 8 | $R_2C_2 + 2\,cAMP \xrightleftharpoons[kb_8]{kf_8} R_2C_2{\cdot}cAMP_2$ | mass action | $R_8 = k_{f8}[R_2C_2][cAMP]^2 - k_{b8}[R_2C_2cAMP_2]$ |
| 9 | $R_2C_2{\cdot}cAMP_2 + 2\,cAMP \xrightleftharpoons[kb_9]{kf_9} R_2C_2{\cdot}cAMP_4$ | mass action | $R_9 = k_{f9}[R_2C_2cAMP_2][cAMP]^2 - k_{b9}[R_2C_2cAMP_4]$ |
| 10 | $R_2C_2{\cdot}cAMP_4 \xrightleftharpoons[kb_{10}]{kf_{10}} 2\,PKAc + R_2{\cdot}cAMP_4$ | mass action | $R_{10} = k_{f10}[R_2C_2cAMP_4] - k_{b10}[PKAc][R_2cAMP_4]$ |
| Module 5: | PDE Phosphorylation By PKA | | |
| 11 | $PDE1 + \mathbf{PKAc} \xrightarrow{Kcat_{11}} PDE1P$ | Michaelis-Menten | $R_{11} = \frac{K_{cat11}[PDE1][PKAc]}{K_{m11}+[PDE1]} - K_{r11}[PDE1P]$ |
| 12 | $PDE4 + \mathbf{PKAc} \xrightarrow{Kcat_{12}} PDE4P$ | Michaelis-Menten | $R_{12} = \frac{K_{cat12}[PDE4][PKAc]}{K_{m12}+[PDE4]} - K_{r12}[PDE4P]$ |
| Module 6: | cAMP Inhibition | | |
| 13 | $cAMP + \mathbf{PDE1{\cdot}Ca_4{\cdot}CaM} \xrightarrow{Kcat_{13}} AMP$ | Michaelis-Menten | $R_{13} = \frac{K_{cat13}[PDE1Ca_4CaM][cAMP]}{K_{m13}+[cAMP]} - K_{r13}[AMP]$ |
| 14 | $cAMP + \mathbf{PDE4} \xrightarrow{Kcat_{11}} AMP$ | Michaelis-Menten | $R_{14} = \frac{K_{cat14}[PDE4][cAMP]}{K_{m14}+[cAMP]} - K_{r14}[AMP]$ |
| 15 | $cAMP + \mathbf{PDE4P} \xrightarrow{Kcat_{15}} AMP$ | Michaelis-Menten | $R_{15} = \frac{K_{cat15}[PDE4P][cAMP]}{K_{m15}+[cAMP]} - K_{r15}[AMP]$ |

*Enzymes are in bold type

### S2 Table of Parameters

TableS 2: Reaction parameters calculated for the model

| **Reaction rate** | $k_f$ | unit | $k_b$ | unit | $K_{cat}$ | unit | $K_m$ | unit | $K_r$ | unit |
|---|---|---|---|---|---|---|---|---|---|---|
| $R_1$ | 0.1 | $\mu M^{-2} \cdot s^{-1}$ | 1 | $s^{-1}$ | - | - | - | - | - | - |
| $R_2$ | 0.005 | $\mu M^{-2} \cdot s^{-1}$ | 1 | $s^{-1}$ | - | - | - | - | - | - |
| $R_3$ | 45.46 | $\mu M^{-1} \cdot s^{-1}$ | 1 | - | - | - | - | - | - | - |
| $R_4$ | 6.89 | $s^{-1}$ | 0.39 | $s^{-1}$ | - | - | 93.02 | $\mu M$ | - | - |
| $R_5$ | 87.65 | $\mu M^{-1} \cdot s^{-1}$ | 1 | $s^{-1}$ | - | - | - | - | - | - |
| $R_6$ | 49.80 | $s^{-1}$ | 1.09 | $s^{-1}$ | - | - | 6.95 | $\mu M$ | - | - |
| $R_7$ | - | - | - | - | 7.41 | $s^{-1}$ | 193.73 | $\mu M$ | 0.44 | $s^{-1}$ |
| $R_8$ | 0.19 | $\mu M^{-1} s^{-1}$ | 1 | $s^{-1}$ | - | - | - | - | - | - |
| $R_9$ | 27.89 | $\mu M^{-1} s^{-1}$ | 1 | $s^{-1}$ | - | - | - | - | - | - |
| $R_{10}$ | 0.21 | $\mu M^{-1} s^{-1}$ | 1 | $s^{-1}$ | - | - | - | - | - | - |
| $R_{11}$ | - | - | - | - | 0.10 | $s^{-1}$ | 0.42 | $\mu M$ | 0.12 | $s^{-1}$ |
| $R_{12}$ | - | - | - | - | 7.67 | $s^{-1}$ | 1.34 | $\mu M$ | 0.02 | $s^{-1}$ |
| $R_{13}$ | - | - | - | - | 1* | $s^{-1}$ | 35* | $\mu M$ | 0 | $s^{-1}$ |
| $R_{14}$ | - | - | - | - | 8.66 | $s^{-1}$ | 1.21 | $\mu M$ | 0.10 | $s^{-1}$ |
| $R_{15}$ | - | - | - | - | 1.59 | $s^{-1}$ | 0.84 | $\mu M$ | 0 | $s^{-1}$ |

*All the kinetic parameters are calculated by fitting the experimental data to the kinetic equations. For reaction #13, the kinetic parameters are directly extracted from [1].

### S3 Table of Initial Conditions

TableS 3: Initial conditions and diffusion coefficients of species in the model

| No. | Species | Compartment | Initial Concentration | Diffusion Coefficient |
|---|---|---|---|---|
| 1 | $Ca^{2+}$ | Cytosol | 1 $\mu M$* | 174.3 $\frac{\mu m^2}{s}$ [2] |
| 2 | CaM | Cytosol | 1.24 $\mu M$* | 11 $\frac{\mu m^2}{s}$ [2] |
| 3 | $Ca_2$·CaM | Cytosol | 0 | 11 $\frac{\mu m^2}{s}$ [2] |
| 4 | $Ca_4$·CaM | Cytosol | 0 | 11 $\frac{\mu m^2}{s}$ [2] |
| 5 | AC1 | Membrane | 5 $\mu M * \cdot \frac{Volume}{Surface}$ | 0.01 $\frac{\mu m^2}{s}$ † |
| 6 | AC1$Ca_2$·CaM | Membrane | 0 | 0.01 $\frac{\mu m^2}{s}$ † |
| 7 | AC1·CaM | Membrane | 0 | 0.01 $\frac{\mu m^2}{s}$ † |
| 8 | PDE1 | Cytosol | 3.80 $\mu M$* | 5 $\frac{\mu m^2}{s}$ † |
| 9 | PDE1$Ca_2$·CaM | Cytosol | 0 | 5 $\frac{\mu m^2}{s}$ † |
| 10 | PDE1·CaM | Cytosol | 0 | 5 $\frac{\mu m^2}{s}$ † |
| 11 | PDE1P | Cytosol | 0 | 5 $\frac{\mu m^2}{s}$ † |
| 12 | PDE4 | Cytosol | 1 | 5 $\frac{\mu m^2}{s}$ † |
| 13 | PDE4P | Cytosol | 0 | 5 $\frac{\mu m^2}{s}$ † |
| 14 | ATP | Cytosol | 2000 $\mu M$* | 74.7 $\frac{\mu m^2}{s}$ [2] |
| 15 | cAMP | Cytosol | 0 | 86.4 $\frac{\mu m^2}{s}$ [2] |
| 16 | AMP | Cytosol | 0 | 85.5 $\frac{\mu m^2}{s}$ [2] |
| 17 | $R_2C_2$ | Cytosol | 0.22 $\mu M$* | 10 $\frac{\mu m^2}{s}$ † |
| 18 | $R_2C_2$·$cAMP_2$ | Cytosol | 0 | 10 $\frac{\mu m^2}{s}$ † |
| 19 | $R_2C_2$·$cAMP_4$ | Cytosol | 0 | 10 $\frac{\mu m^2}{s}$ † |
| 20 | $R_2$·$cAMP_4$ | Cytosol | 0 | 10 $\frac{\mu m^2}{s}$ † |
| 21 | PKAc | Cytosol | 0 | 8.1 $\frac{\mu m^2}{s}$ [2] |

* Estimated values obtained by fitting the experimental data to the kinetic equations.

† Generic values.

#### S4 Input Calcium Function

For the 3D spatial model, the input calcium is modeled as a function with sinusoidal oscillations with exponential decay. Every five minutes, there is a calcium burst and inside of each calcium burst there are calcium pulses with 0.5 Hz oscillations:

Input $Ca^{2+}$ Function $= (0.45\sin(\pi(t-0.5)/1)+0.45)(0.505(1-0.01)^{t})(1+\tanh(\pi(t+14.543)/10))+0.1$

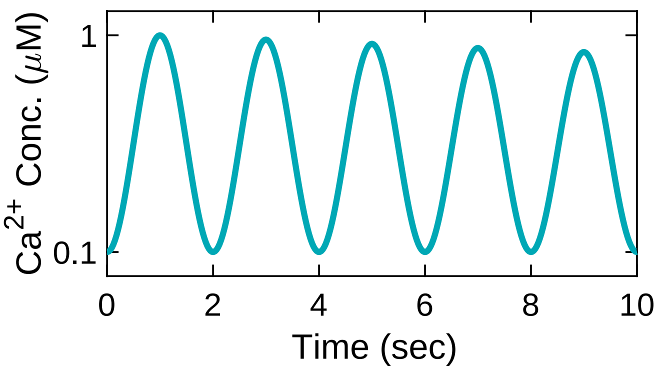


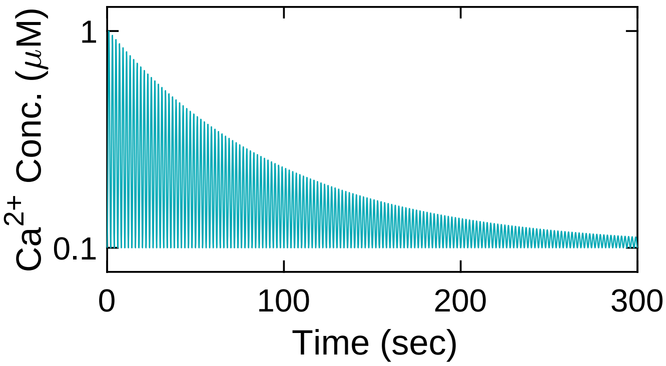


FigureS 1: Input calcium for the spatial model with 0.5 Hz pulses inside every 300-second bursts.

#### S5 ATP and cAMP Oscillations in the Spatial Model

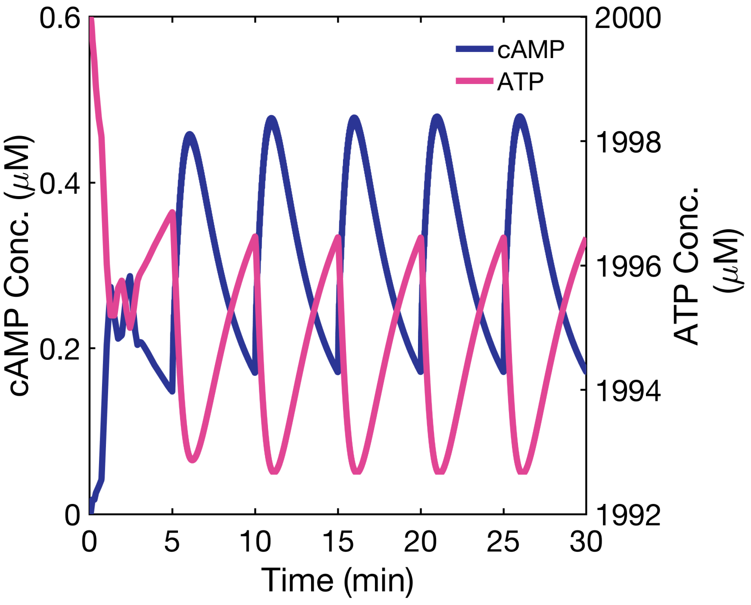


FigureS 2: ATP Oscillating with cAMP in the spatial model. Note that the change in ATP concentration is minimal.

#### S6 Effect of Calcium Diffusion Coefficient on the cAMP Dynamics

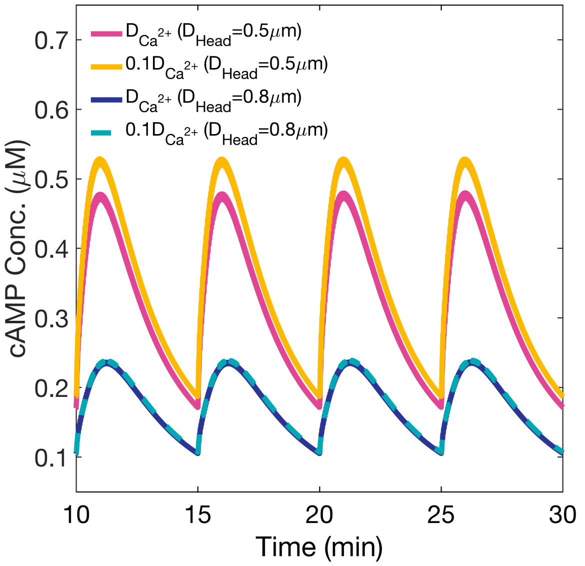


FigureS 3: Effect of the calcium diffusion coefficient on cAMP dynamics in the small and large spherical heads. Note that the diffusion coefficient does not change the spatiotemporal dynamics of cAMP in our model.
